# Modeling cross-regulatory influences on monolignol transcripts and proteins under single and combinatorial gene knockdowns in *Populus trichocarpa*

**DOI:** 10.1101/677047

**Authors:** Megan L. Matthews, Jack P. Wang, Ronald Sederoff, Vincent L. Chiang, Cranos M. Williams

**Affiliations:** Department of Electrical and Computer Engineering, North Carolina State University, Raleigh, NC, USA; State Key Laboratory of Tree Genetics and Breeding, Northeast Forestry University, Harbin, China; Department of Forestry and Environmental Resources, Forest Biotechnology Group, North Carolina State University, Raleigh, NC, USA; Department of Forest Biomaterials, North Carolina State University, Raleigh, NC, USA

## Abstract

Accurate manipulation of metabolites in the monolignol biosynthetic pathway is a key step for controlling lignin content, structure, and other wood properties important to the bioenergy and biomaterial industries. A crucial component of this strategy is predicting how single and combinatorial knockdowns of monolignol specific gene transcripts influence the abundance of monolignol proteins, which are the driving mechanisms of monolignol biosynthesis. Computational models have been developed to estimate protein abundances from transcript perturbations of monolignol specific genes. The accuracy of these models, however, is hindered by the inability to capture indirect regulatory influences on other pathway genes. Here, we examine the manifestation of these indirect influences collectively on transgenic transcript and protein abundances, identifying putative indirect regulatory influences that occur when one or more specific monolignol pathway genes are perturbed. We created a computational model using sparse maximum likelihood to estimate the resulting monolignol transcript and protein abundances in transgenic *Populus trichocarpa* based on desired single or combinatorial knockdowns of specific monolignol genes. Using *in-silico* simulations of this model and root mean square error, we show that our model more accurately estimates transcript and protein abundances in differentiating xylem tissue when individual and families of monolignol genes were perturbed. This approach provides a useful computational tool for exploring the cascaded impact of single and combinatorial modifications of monolignol specific genes on lignin and other wood properties. Additionally, these results can be used to guide future experiments to elucidate the mechanisms responsible for the indirect influences.

**Author summary:** Engineering trees to have desirable lignin and wood traits is of significant interest to the bioenergy and biomaterial industries. Genetically modifying the expression of the genes that drive the monolignol biosynthetic pathway is a useful method for obtaining new traits. Modifying the expression of one gene affects not only the abundance of its encoded protein, but can also indirectly impact the amount of other transcripts and proteins. These proteins drive the monolignol biosynthetic pathway. Having an accurate representation of their abundances is key to understanding how lignin and wood traits are altered. We developed a computational model to estimate how the abundance of monolignol transcripts and proteins are changed when one or more monolignol genes are knocked down. Specifying only the abundances of the targeted genes as input, our model estimates how the levels of the other, untargeted, transcripts and proteins are altered. Our model captures indirect regulatory influences at the transcript and protein levels observed in experimental data. The model is an important addition to current models of lignin biosynthesis. By incorporating our approach into the existing models, we expect to improve our ability to explore how new combinations of gene knockdowns impact lignin and many other wood properties.

## Introduction

Lignin is an important phenylpropanoid polymer that is embedded with cellulose and hemicelluloses in plant secondary cell walls [1, 2]. It plays an important role in plant physiology, defense, and adaptation by providing structural integrity, conducting water through vascular tissues, and acting as a barrier to pests and pathogens [1, 3]. Lignin is composed of three main sub-units, the *p*-hydroxyphenyl (H), guaiacyl (G), and syringyl (S) monolignols. These monolignols define the composition and interunit linkages that determine other characteristics of lignin [1, 2, 4]. How these monolignols are formed and synthesized into lignin has been an important research area for more than five decades [5]. Producing plants that have specific lignin phenotypes is of significant interest in the bioenergy and biomaterial industries [6, 7].

A key step to controlling lignin phenotypes is by precise manipulation of the monolignol biosynthesis pathway. Genetic modifications are a useful method for manipulating metabolic pathway behavior. These modifications alter transcript production or abundance resulting in a change to the amount of proteins available to catalyze key pathway reactions. It is not always intuitive how genetic modifications propagate through biological systems culminating in changes to phenotypic traits. Many approaches have been presented to understand phenotypic changes based on single layers of biological information, such as GWAS [8, 9]. However, biological systems regulate themselves through diverse mechanisms including, transcriptional [10–12] and post-transcriptional [10, 13, 14] regulation, and post-translational modifications [14–16] among others. By improving our understanding of the factors that arise when knocking down genes, we can better discern how metabolic pathway activity and phenotypic responses change in response to knockdowns and other modifications.

Extensive study of the metabolic reactions associated with monolignol biosynthesis in *P. trichocarpa* has resulted in a detailed mechanistic computational model of the pathway, composed of 24 ordinary differential equations with 104 Michaelis-Menten and 103 inhibition kinetic parameters [17, 18]. Wang et al. expanded their mechanistic metabolic model of the monolignol pathway to incorporate information spanning the genome, transcriptome, proteome, and 25 lignin and wood traits [4]. This multi-scale model was used to help identify novel combinatorial genetic modifications that result in desired lignin and wood characteristics such as increased saccharification efficiency without negatively impacting plant growth. Wang et al., made the simplifying assumption that the abundance of each protein was dependent only on the transcript abundance of its monolignol gene. This simplification ignores possible epistatic regulatory interactions that exist among the monolignol gene transcripts and proteins.

Regulatory mechanisms can act at many different points in biological pathways. The most commonly studied are transcriptional regulatory networks. Inferring and modeling the relationships in these networks has been an area of significant interest [19–24]. Temporal measurements of transcript abundance in response to stress or gene perturbations are often used to identify and model these networks [21, 23]. In the absense of temporal data, variations in steady-state transcript abundance from multiple gene perturbation experiments [25, 26] or naturally occuring variations characterized using eQTLs [27–29] have been used. Network structure and parameter estimation are the two main components of modeling regulatory networks. These steps are often combined using a regularization term to penalize model complexity during parameter estimation [20]. Examples of these approaches include LASSO [27, 30, 31], LARS [32–34], and sparse maximum likelihood [29].

In addition to transcriptional regulation, post-transcriptional and post-translational regulatory mechanisms relating to translation or protein degradation can play a critical role in protein abundance [13, 35, 36]. Differences in expression of transcripts and proteins suggesting such regulatory mechanisms have been found for genes encoding cell wall proteins in *Arabidopsis thaliana* [37, 38] and for genes involved in tobacco xylem cell differentiation [39]. These mechanisms were also proposed to explain the poor correlations between some of the monologinol gene transcripts and proteins in some transgenic *P. trichocarpa* [4]. Having an accurate representation of the protein abundance profile is important to assess how the metabolic pathway is driven. Developing a computational model that captures the indirect regulatory influences between monolignol genes at both the transcript and protein levels is important for exploring how novel transgenic modifications impact lignin and wood characteristics.

In this paper we perform differential abundance analyses on the monolignol gene transcript and protein abundances to further characterize epistatic influences on the expression of the monolignol genes in differentiating xylem tissue of *P. trichocarpa*. We then used the experimental transcript and protein abundance measurements [4] to develop a model that describes the indirect relationships between the monolignol genes as transcript to transcript, transcript to protein, protein to transcript, and protein to protein influences. We used a *sparse maximum likelihood* estimator [29] to identify potential key indirect regulatory influences between the monolignol gene transcripts and proteins. Through *in-silico* simulations, our model more accurately estimates monolignol transcript and protein abundances in transgenic plants where individual and families of monolignol genes were knocked down than a model that does not incorporate such regulatory influences. We identified and modeled apparent regulatory influences among the *PtrCAld5H*, *Ptr4CL*, *PtrPAL*, *PtrC3H3*, *PtrC4H*, and *PtrHCT* gene families and among the *PtrHCT*, *Ptr4CL* gene families and *PtrCCoAOMT3*, which manifest as relationships between protein abundances but not the transcripts. Our model is able to capture many of these putative epistatic influences between the monolignol transcripts and proteins by specifying the abundance level of the targeted transcript as an input. This model provides an important addition to the current computational lignin model, allowing for the further exploration of the cascaded impact of genetic modifications on the content, composition, and interunit structure of lignin and its related wood properties. The identified relationships can also be used to further investigate the specific regulatory mechanisms that govern monolignol gene expression.

## Results

### Data description

Wang et al. [4] performed a series of systematic transgenic experiments that knocked down each of the 21 lignin specific genes and their gene families in the model tree *P. trichocarpa*. The absolute transcript abundances were measured using RNAseq, and the absolute protein abundances were obtained using protein cleavage coupled with isotope dilution mass spectrometry (PC-IDMS) [40]. Multiple independent lines were grown for each transgenic construct. Usually three of those lines were selected to show the effects of a range in the level of the targeted knockdown gene expression, providing an indication of the complexity of putative interactions as responses can be linear or nonlinear. For each line, up to three biological replicates were collected after six months of growth, resulting in 207 transgenic measurement profiles and 18 wildtype measurement profiles. Due to limited greenhouse space, these experiments were grown in six batches. To account for batch effects on the data, Wang et al. normalized the data to the wildtype mean in each batch [4]. Additionally, the PC-IDMS approach for quantifying protein abundance was not able to differentiate between the *PtrPAL4* and *PtrPAL5* proteins because of the near identity of these proteins [40]. The transcript and protein abundances for *PtrPAL4* and *PtrPAL5* were combined into one, which we refer to as *PtrPAL4/5*.

### Differential abundance analysis

To further examine the influence of targeted knockdowns on other non-targeted genes, we performed a differential abundance analysis on both the transcripts and protein data. Fig 1 contains heatmaps showing the results for five of the knockdown experiments: construct i69, which targeted *PtrC3H3*, *PtrC4H1*, and *PtrC4H2* (Fig 1A); construct i29, which targeted *PtrCAld5H1* and *PtrCAld5H2* (Fig 1B); construct i35, which targeted *PtrCAD1* and *PtrCAD2* (Fig 1C); construct i15, which targeted *Ptr4CL3* and *Ptr4CL5* (Fig 1D); and construct i21, which targeted *PtrCCoAOMT3* (Fig 1E). Heatmaps for the remaining transgenics can be found in Supplemental Figs S1-S4. Each column represents a different line of that experiment, with each line containing up to 3 replicates. The rows indicate the monolignol specific gene name with the purple names indicating the gene(s) that were knocked down. The colorscale of these heatmaps corresponds to the log fold change (logFC) from their wildtype. Red represents a negative fold change, i.e., a decrease in expression, and green corresponds to a positive fold change or an increase in expression. Gray boxes represent missing data. Changes in abundance that had a *p*-value adjusted for multiple comparisons less than 0.05 are considered statistically significant and are indicated with an asterisk.

**Fig 1.**
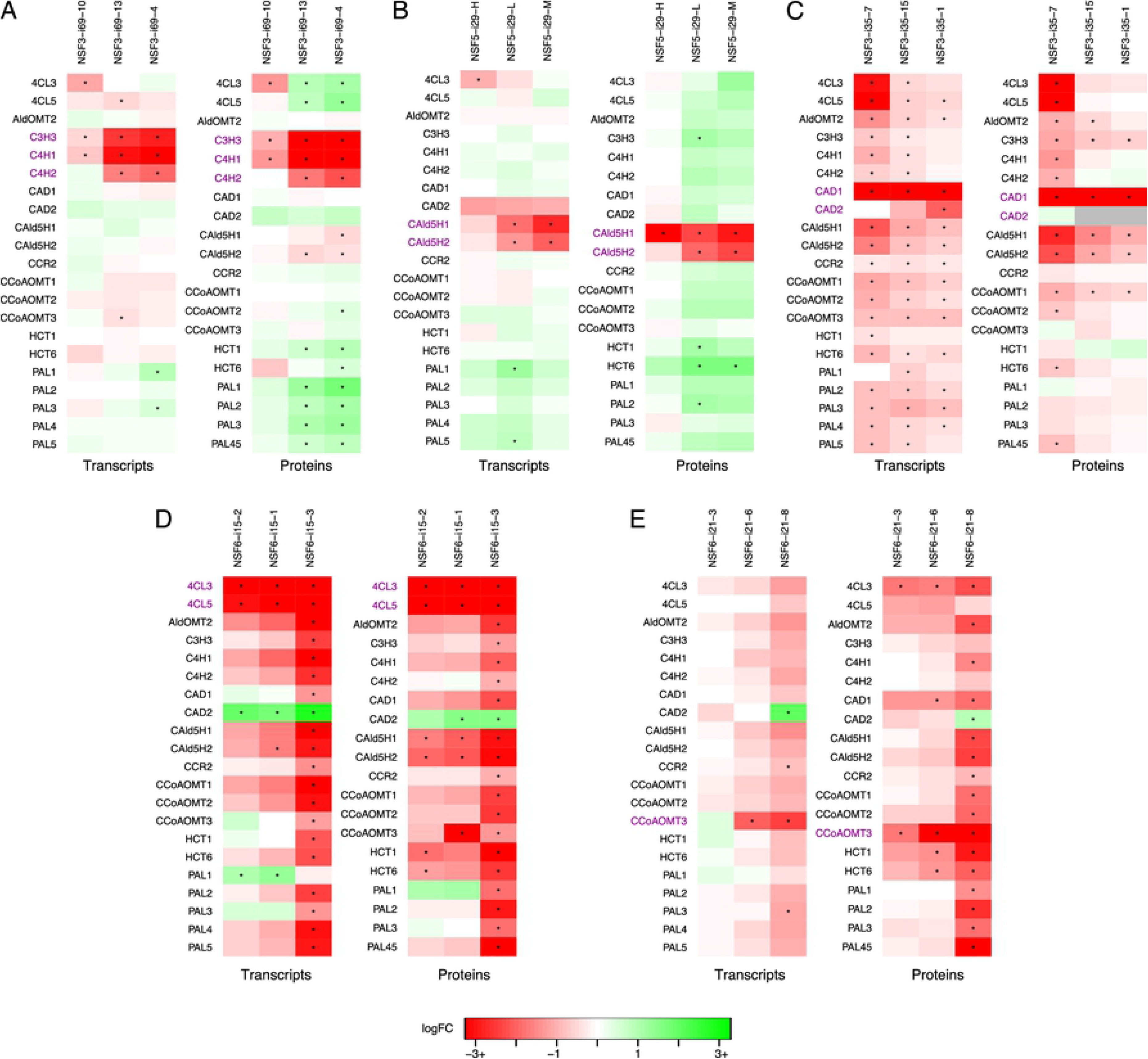
Monolignol gene transcript and protein differential abundance. (A) *PtrC3H3*, *PtrC4H1* and *PtrC4H2* knockdown experiments (Construct i69). (B) *PtrCAld5H1* and *PtrCAld5H2* knockdown experiments (Construct i29). (C) *PtrCAD1* and *PtrCAD2* knockdown experiments (Construct i35). (D) *Ptr4CL3* and *Ptr4CL5* knockdown experiments (Construct i15). (E) *PtrCCoAOMT3* knockdown experiments (Construct i21). Gray boxes are due to missing data. Rows are the monolignol gene names, with the targeted genes for each experiment in purple. Columns are the experimental lines. ∗ indicates *p*_adj_<0.05.

We see significant changes in abundance in several of the untargeted monolignol genes. This indicates that there are cross-influences among the targeted monolignol genes impacting the abundances of untargeted monolignol transcripts and proteins. Collectively examining the responses of both the monolignol gene transcripts and proteins provides insight to the regulatory influences between the monolignol genes that would not be detected by just examining the transcripts. While we observe some instances of the same differential abundance patterns in the transcripts and proteins, suggesting transcriptional regulation, we also observe several cases where only a monolignol gene’s transcript or its protein abundance is significantly altered. This suggests the presence of post-transcriptional or post-translational regulation.

In the *PtrC3H3*, *PtrC4H1*, and *PtrC4H2* knockdown experiments we observe significant increases in the abundances of the *Ptr4CL*, *PtrHCT*, and *PtrPAL* proteins and significant decreases in the *PtrCAld5H* proteins (Fig 1A). However, their corresponding transcript abundances, with the exception of some of the *PtrPAL* transcripts, are not found to be differentially expressed. Similarly, in the *PtrCAld5H1* and *PtrCAld5H2* knockdown experiments we observe significant increases in the abundances of the *PtrHCT* and *PtrC3H3* proteins that are not observed in the transcript data (Fig 1B). In the *PtrCAD1* and *PtrCAD2* knockdown experiments (Fig 1C), we observe a decrease in the abundance of both the transcripts and proteins of *PtrCAld5H1* and *PtrCAld5H2*, as well as most of the other monolignol transcripts. Despite this, many of the proteins are not significantly different from their wildtype levels. This could be explained by the same behavior as in the *PtrCAld5H1* and *PtrCAld5H2* knockdowns and *PtrC3H3*, *PtrC4H1*, and *PtrC4H2* knockdown experiments where we also observed an increase in several of the protein abundances. The increase we observe in the proteins in those two knockdowns could lead to wildtype levels in the *PtrCAD1* and *PtrCAD2* knockdown experiments because the transcript abundances are significantly decreased. This behavior is seen to a lesser degree in the experimental line that had the largest decrease in the *Ptr4CL3* and *Ptr4CL5* transcripts and proteins. Additionally, we do not observe this behavior in the *Ptr4CL3* and *Ptr4CL5* knockdown experiments (Fig 1D), suggesting that large knockdowns of the *Ptr4CL* gene family may trump other regulatory influences.

In the *Ptr4CL3* and *Ptr4CL5* transgenics (Fig 1D), we observe significant decreases in abundance of both the transcripts and proteins of *PtrCAld5H1* and *PtrCAld5H2* and an increase in the *PtrCAD2* abundances across multiple transgenic lines. Significant decreases in abundance are also observed in the *PtrHCT1*, *PtrHCT6*, and *PtrCCoAOMT3* proteins in multiple lines. Similar behavior is seen in the transgenics that individually knocked down *Ptr4CL3* (Fig S4A) and *Ptr4CL5* (Fig S4B), with significant decreases observed in the *PtrHCT1*, *PtrHCT6*, *PtrCCoAOMT3*, and *PtrCAD1* proteins. The *PtrHCT1*, *PtrHCT6*, *Ptr4CL3*, and *PtrCAD1* proteins are also significantly decreased in the *PtrCCoAOMT3* transgenics (Fig 1E). There are multiple transgenics where one line showed significant changes in all or almost all of the monolignol transcripts and proteins, but not in the other lines for the same transgenic such as i35-7 (Fig 1C), i15-3 (Fig 1D), i19-7 (Fig S3F), and a13-6 (Fig S4B). This behavior could be due to a nonlinear response to a change in the abundance of one or more of the monolignol transcripts and proteins.

Some of the observed indirect effects occur within gene families, such as in the *PtrPAL* knockdowns (Figs S1A-D), the *PtrCCoAOMT1* knockdowns (Fig S3C), the *PtrCAld5H1* and *PtrCAld5H2* single knockdowns (Figs S3D and E), and in the *Ptr4CL3* and *Ptr4CL5* single knockdowns (Figs S4A and B). These indirect effects within gene families could be due to sequence relationships with the targeted gene instead of regulatory mechanisms.

Capturing the effect of these indirect regulatory influences is necessary to effectively estimate the resulting protein levels that are responsible for driving monolignol biosynthesis. Further, it is necesarry to capture the indirect effects that affect the transcripts and the indirect effects on the proteins separately.

### Computational model

We developed a computational model that describes the observed cross-talk or interactions among the monolignol genes by representing each monolignol transcript and protein as a linear combination of the other monolignol transcripts and proteins. This formulation allows us to describe the indirect cross-influences as transcript to transcript and protein to transcript influences to represent influences impacting transcription, and transcript to protein and protein to protein influences to represent the indirect influences affecting the protein abundances. We estimated the weights of the connections that make up these linear combinations using a sparse maximum likelihood algorithm and the mean abundances from the experimental lines (see Methods and S1 Text). Using this model, we simulated the response of the untargeted monolignol gene transcripts and proteins based on the desired transcript abundance of a targeted monolignol gene or gene family (Fig 2A). We compare our model with the model from Wang et al. [4] which assumed that all of the protein abundances were proportional to their transcript levels (Fig 2B). We compare our model to two specific scenarios of this old model: scenario 1, where the desired targeted transcript levels are specified and the untargeted transcripts remain at wildtype levels, and scenario 2 where the full transcript profile is specified. We estimate the untargeted monolignol transcript and protein abundances using our model and both scenarios of the old model for single gene and gene family knockdowns corresponding to the transgenic experiments [4]. When exploring novel combinatorial knockdowns, however, where complete transcript profiles are unknown, scenario 2 cannot be simulated. We refer to the transcript of a gene as tGENE and the protein of a gene as pGENE in the following sections.

**Fig 2.**
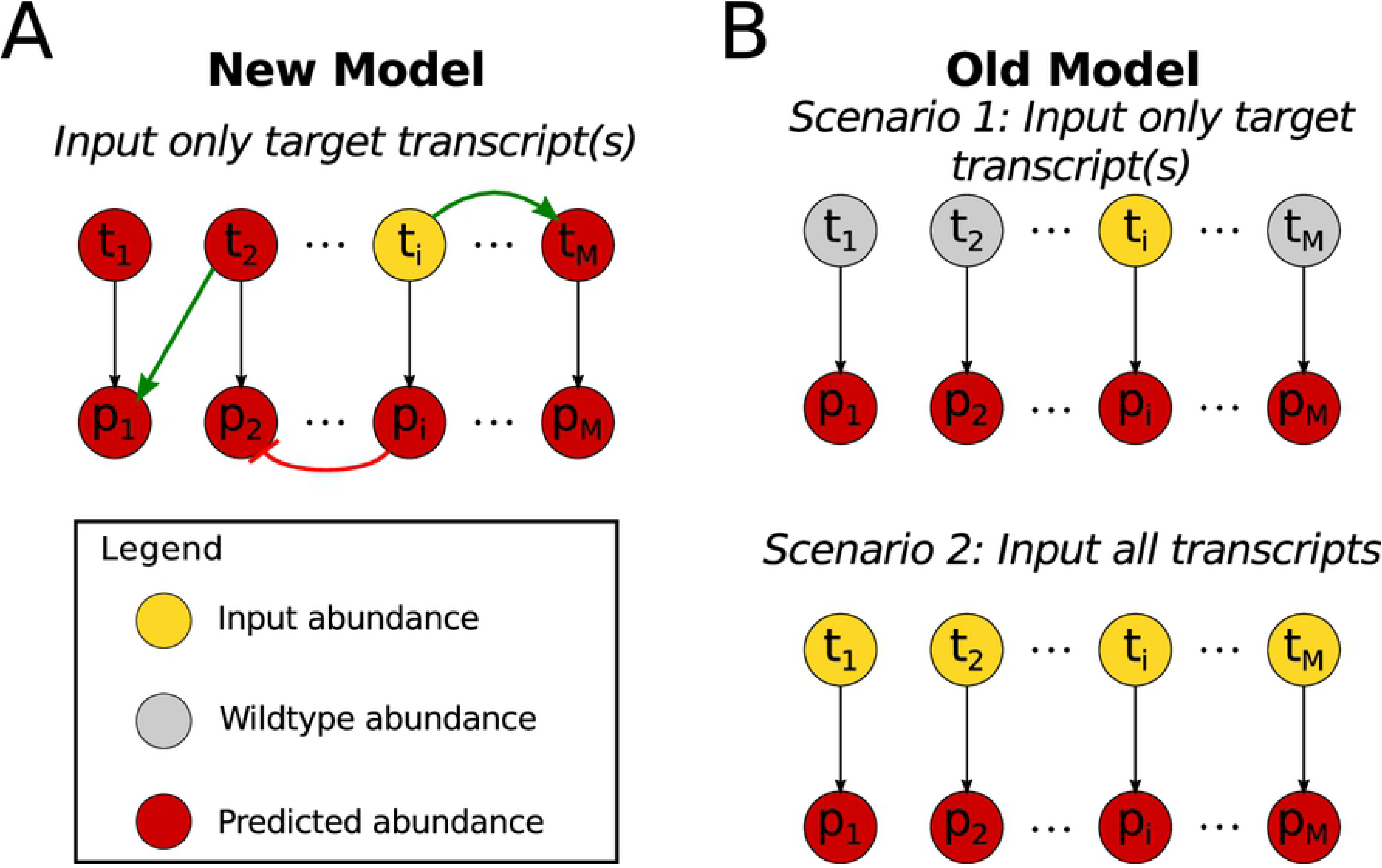
Diagram of transcript-protein models. (A) Diagram describing our model which includes positive (green arrows) and negative (red arrows) influences among the monolignol transcripts and proteins defined by **B** (Eq. (3)). Using only targeted input abundances (yellow), the other untargeted monolignol transcripts and proteins are predicted (red) (B) In the old model only the one-to-one relationships from a monolignol transcript to its protein were included. In scenario 1, only the targeted monolignol transcripts were used as input abundances (yellow), the untargeted transcripts remained at wildtype levels (gray) and the protein abundances were predicted (red). In scenario 2, all of the monolignol transcript abundances were used as input (yellow) to predict (red) the monolignol protein abundances.

We performed a 10×10-fold cross-validation resulting in 100 training and testing folds. The proposed model and the old model were trained on each of the 100 training folds. For each of the trained models, the knockdown experiments in the training fold and corresponding testing fold were emulated following the model estimation procedure (see Methods) for our model, and following scenario 1 for the old model. In each of these emulated experiments, the trained models estimated the untargeted monolignol gene transcripts and proteins. Fig 3 shows boxplots of the resulting root mean square errors (RMSE) of the estimated abundances across the 100 training (Fig 3A and 3C) and 100 testing folds (Fig 3B and 3D) for both our proposed model and the old model (Fig 2A and B - scenario 1). We performed a t-test to compare the distributions of the RMSEs from the new model and the old model for each monolignol transcript and protein. The x-axis labels with an asterisk had a significant difference (*p*<0.05) in the means of the distributions from the new model (red) and scenario 1 of the old model (yellow). We see that all of the training sets were shown to have a significant difference (Figs 3A and 3C) while in the testing sets, 14 out of 20 of the transcripts and 11 out of 20 of the proteins were shown to have a significant difference (Figs 3B and 3D). In each of the significant cases, the distributions from the new model have a lower mean RMSE.

**Fig 3.**
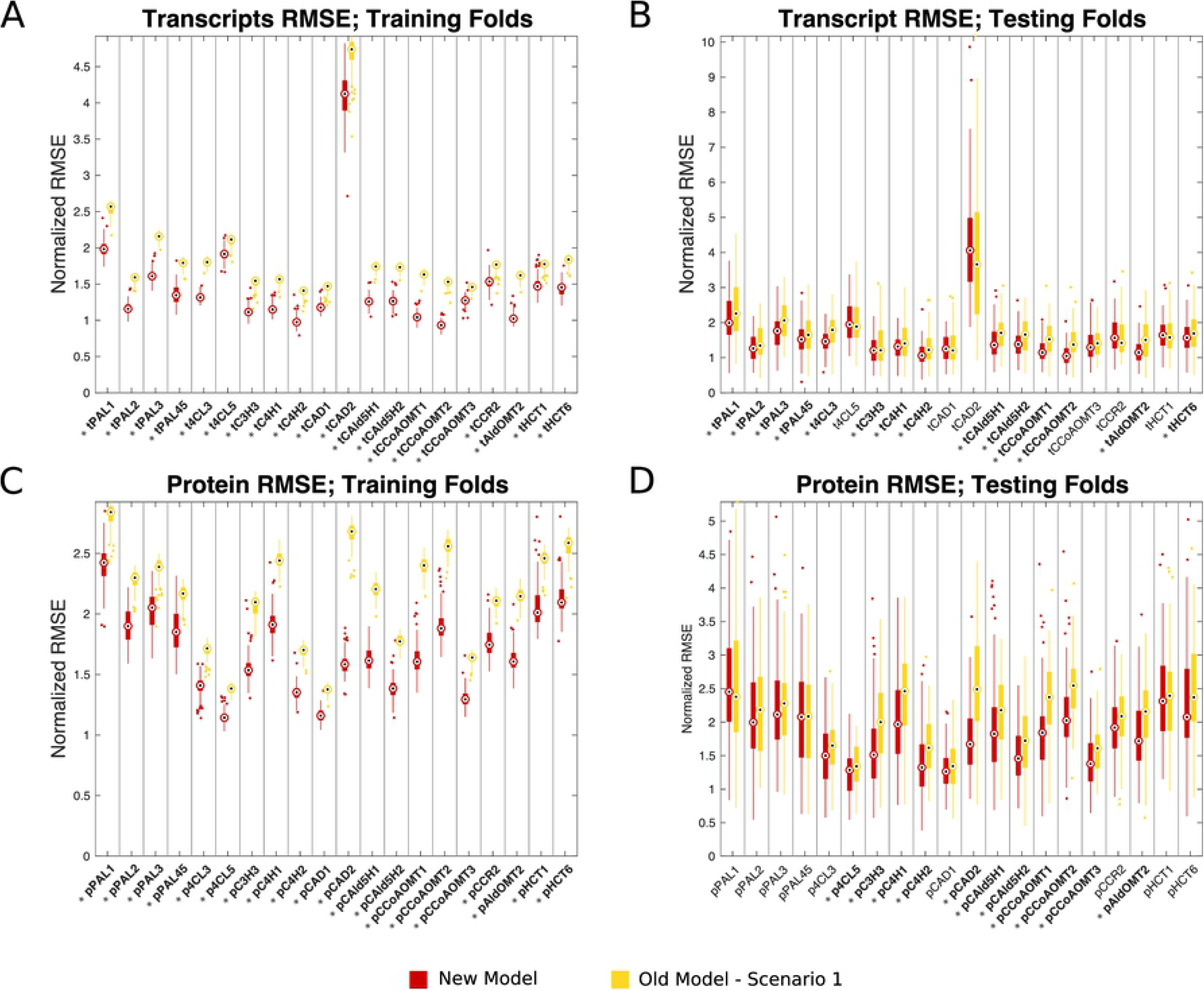
Boxplots of the RMSEs from the 10×10-fold cross-validation. Central marks indicate the medians and the bottom and top edges of each box indicate the 25th and 75th percentiles respectively. For these plots we normalized the RMSE for each monolignol transcript and protein by their corresponding standard deviations estimated from the wildtype experiments. This normalization allows each of the monolignol transcripts and proteins to be viewed on similar scales. Since the RMSEs from both models are scaled the same, this does not alter the interpretation of the results. Red boxes are from our new model, and the yellow boxes are from the old model. (A) Transcripts: training folds. (B) Transcripts: testing folds. (C) Proteins: training folds. (D) Proteins: testing folds.

These cross-validation results show that our model performs as well or better than the scenario 1 of the old model.

Fig 4A shows a heatmap of the relationships identified in our model (**B** in Eq 3) when trained on the means from all of the experimental lines. Green represents a positive influence, and red represents a negative influence. Each column represents the transcript or protein that is the source of an influence, and the row represents the transcript or protein that is being influenced. The top left quadrant contains the transcript to transcript influences, the top right quadrant contains the protein to transcript influences, the bottom left quadrant contains the transcript to protein influences, and the bottom right quadrant contains the protein to protein influences. There were 295 relationships detected out of a possible 1540 (19.16% sparse). The full set of relationships and their weights for our model can be found in Table S1. For comparison, Fig 4B shows the equivalent representation of the old model, which just contains the *t*_*i*_ → *p*_*i*_ relationships. As expected, a positive influence was detected for each transcript to its associated protein (*t*_*i*_ → *p*_*i*_). The transcript to transcript and protein to protein influences make up the majority of the remaining influences estimated. There are not many protein to transcript influences detected, suggesting that protein abundances that are altered due to post-transcriptional or post-translational mechanisms may not result in changes at the transcriptional level that you would see with a targeted knockdown of that gene. Such as when the abundance of pCAD1 is decreased in the *PtrCCoAOMT3* (Fig 1E), *Ptr4CL3* (Fig S4A), or *Ptr4CL5* (Fig S4B) knockdowns, but the changes in transcript abundance that occur when *PtrCAD1* is knocked down (Fig 1C, Fig S2C) are not observed.

**Fig 4.**
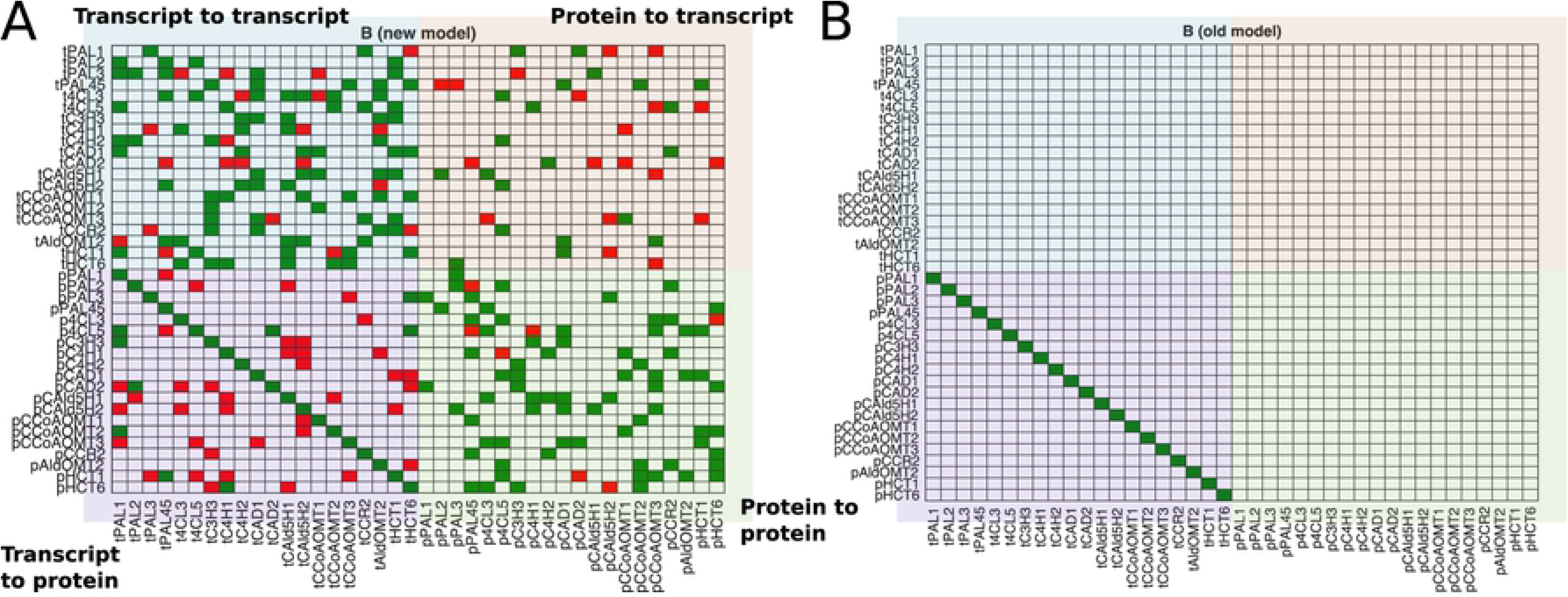
Heatmaps of the relationships in the transcript-protein models. (A) Heatmap of the edge matrix **B** (Eq 3) solved using a sparse maximum likelihood estimator. Green for positive influence, red for negative influence. Edges are from columns to rows (e.g., the first row shows edges tPAL3 → tPAL1, tCCR2 → tPAL1, tHCT6 → tPAL1, pC3H3 → tPAL1, pCAD2 → tPAL1, pCAld5H2 → tPAL1, and pCCoAOMT3 → tPAL1). There were 295 edges detected out of a possible 1540 (19.16% sparse). (B) The corresponding heatmap for the relationships considered in the old model (t_*i*_ → p_*i*_).

To further evaluate how well our model captures these cross-influences affecting the monolignol transcript and protein abundances, we used our model and scenarios 1 and 2 of the old model to emulate the five transgenic experiments from our differential abundance analysis. For each of the five targeted experiments, we further described the results from the models for a subset of the untargeted monolignol genes that had a significant change in the abundance of their transcripts, proteins, or both in the differential abundance analysis.

### *PtrC3H3*, *PtrC4H1*, and *PtrC4H2* knockdowns

Three experimental lines were analyzed where *PtrC3H3*, *PtrC4H1*, and *PtrC4H2* were knocked down (Fig 5). From the differential abundance analysis, there were 5 transcripts and 11 proteins of the untargeted genes that had a significant change in abundance in at least one of the experimental lines, which are signified by an asterisk (Figs 1A, 5A). We include significant changes that occur in at least one of the lines since each line represents a different amount of knockdown of the targeted genes. We selected *Ptr4CL5*, *PtrCAld5H2*, and *PtrHCT1* transcripts and proteins to compare the simulated results from our model with scenarios 1 and 2 of the old model. Fig 5B shows the levels of knockdown for each of the three lines for the *PtrC3H3*, *PtrC4H1*, and *PtrC4H2* transcripts. The knockdowns range from ~65% to ~10% of wildtype levels for tC3H3 and tC4H1, and ~110% to ~25% of wildtype levels for tC4H2. These tC3H3, tC4H1, and tC4H2 abundances were used to emulate these knockdown experiments in our model and scenario 1 of the old model. For scenario 2 of the old model, measurements from all of the monolignol transcripts were used.

**Fig 5.**
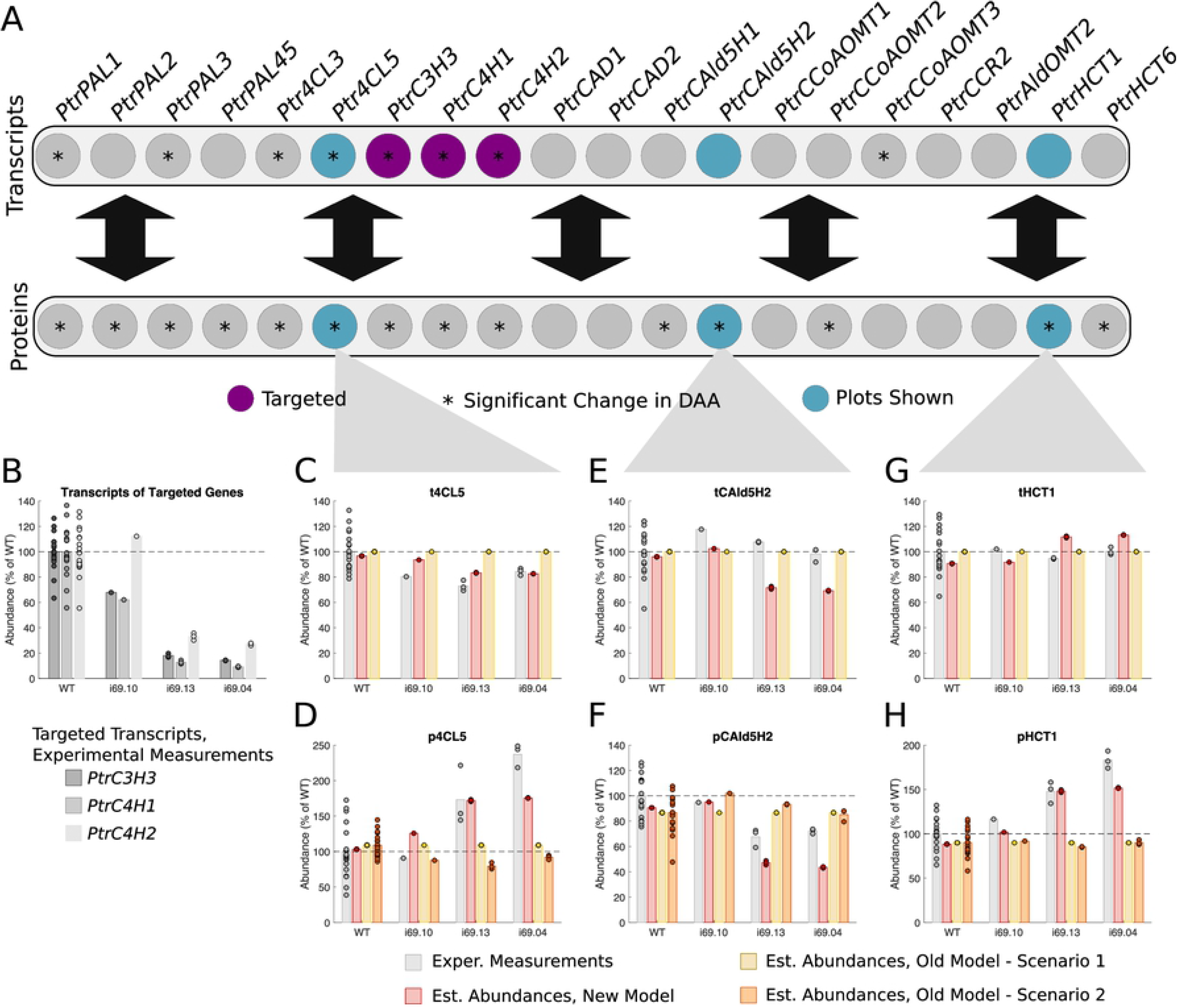
Experimental and estimated abundances of untargeted monolignol gene transcripts and proteins under *PtrC3H3*, *PtrC4H1*, and *PtrC4H2* knockdowns. (A) Diagram showing targeted monolignol gene transcripts (purple), the transcripts and proteins that were found to have a significant change in abundance in at least one of the experimental lines (∗). (B) Level of knockdown of the targeted gene transcripts across the experimental lines. Experimental and estimated untargeted monolignol gene transcript and protein abundances for (C) t4CL5, (D) p4CL5, (E) tCAld5H2, (F) pCAld5H2, (G) tHCT1, and (H) pHCT1.

A slight decrease to ~80% average wildtype levels was experimentally measured for t4CL5 (Fig 5C), but an increase up to ~250% was measured for p4CL5 (Fig 5D). Our model captured this behavior, estimating a decrease in t4CL5 to ~80% of wildtype but an increase in p4CL5 to ~175% of wildtype levels. Neither scenario of the old model captured the increase in p4CL5, with scenario 2 estimating a small decrease in p4CL5 to ~80% corresponding to the decrease measured in t4CL5.

No change from wildtype levels was experimentally measured for tCAld5H2 (Fig 5E). For pCAld5H2 a decrease to ~70% of wildtype levels was experimentally measured (Fig 5F). For both tCAld5H2 and pCAld5H2, our model over-estimated a decrease in the abundances to ~70% of wildtype for tCAld5H2 and ~45% of wildtype for pCAld5H2. While scenario 2 of the old model did not estimate any change in the abundance of pCAld5H2.

For all three lines, the measured abundances for tHCT1 remain around wildtype levels (Fig 5G) while an increase in pHCT1 is experimentally measured ranging up to ~180% of wildtype levels (Fig 5H). Our model estimated wildtype level abundances for tHCT1, and an increase up to ~150% of wildtype levels for pHCT1, which are consistent with the experimentally measured values. Because there was no change in the experimental transcript abundances, scenario 2 of the old model incorrectly estimates no change in the abundance of pHCT1.

Neither scenario of the old model captured the increase in the *Ptr4CL5* and *PtrHCT1* proteins or the decrease in the *PtrCAld5H2* protein. Alternatively, our new model successfully estimated these changes in the three proteins, estimated the decrease in t4CL5 and estimated tHCT1 to remain around wildtype levels. It did, however, predict a slight decrease in tCAld5H2 abundance, which was not observed experimentally.

### *PtrCAld5H1* and *PtrCAld5H2* knockdowns

Three experimental lines were analyzed where *PtrCAld5H1* and *PtrCAld5H2* were targeted, knocking them down to values seen in experimental constructs (Fig 6). From the differential abundance analysis, there were 3 transcripts and 4 proteins of untargeted genes that showed significant changes in abundance in at least one of the experimental lines (Figs 1B, 6A). From these, we selected the *PtrPAL2*, *PtrC3H3*, and *PtrHCT6* transcripts and proteins to compare the simulated results from our model with scenarios 1 and 2 of the old model. Fig 6B shows the levels of knockdown, ranging from ~80% to ~20% of wildtype levels, for each of the three lines for the *PtrCAld5H1* and *PtrCAld5H2* transcripts. These tCAld5H1 and tCAld5H2 abundances were used to emulate these knockdown experiments in our model and scenario 1 of the old model. For scenario 2 of the old model, measurements from all of the monolignol transcripts were used.

**Fig 6.**
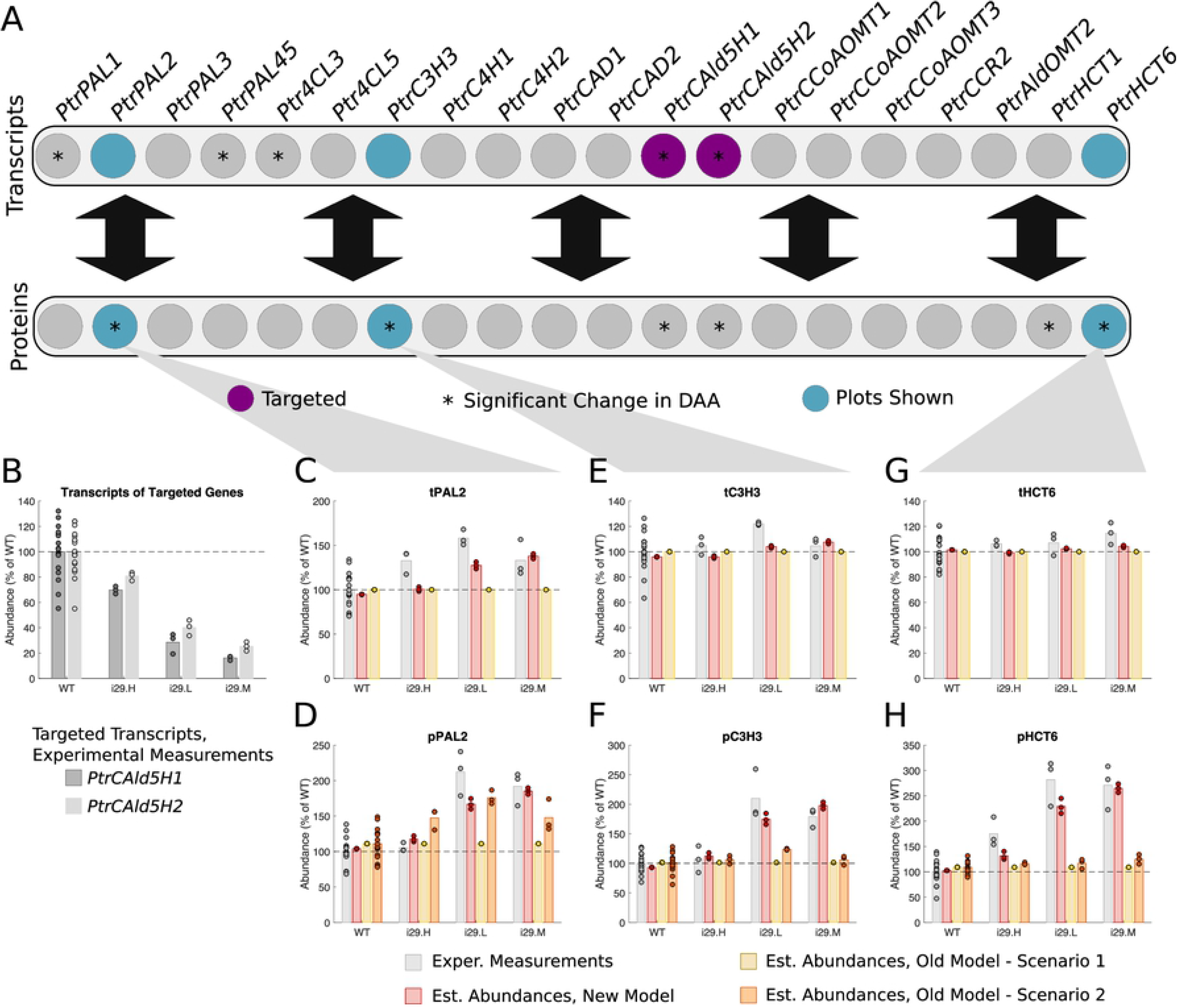
Experimental and estimated abundances of untargeted monolignol gene transcripts and proteins under *PtrCAld5H1* and *PtrCAld5H2* knockdowns. (A) Diagram showing targeted monolignol gene transcripts (purple), the transcripts and proteins that were found to have a significant change in abundance in at least one of the experimental lines (∗). (B) Level of knockdown of the targeted gene transcripts across the experimental lines. Experimental and estimated untargeted monolignol gene transcript and protein abundances for (C) tPAL2, (D) pPAL2, (E) tC3H3, (F) pC3H3, (G) tHCT6, and (H) pHCT6.

Increases up to ~150% of wildtype levels and ~200% of wildtype levels were experimentally measured for tPAL2 (Fig 6C) and pPAL2 (Fig 6D) respectively. Our model estimated increases in abundance up to ~140% of wildtype levels for tPAL2 and ~185% of wildtype levels for pPAL2, which are consistent with the experimentally measured abundances. Scenario 2 of the old model was also consistent with the experimentally measured pPAL2 abundances, estimating an increase up to ~175% of wildtype levels.

Our model estimated wildtype level abundances for tC3H3 which is consistent with the experimentally measured abundances (Fig 6E). For pC3H3 an increase up to ~210% of wildtype levels was experimentally measured (Fig 6)F. Our model was consistent with these results, estimating an increase up to ~200% of wildtype levels. Scenario 2 of the old model, however, was not consistent with the experimental measurements, and estimated wildtype levels for pC3H3.

Our model estimated wildtype level abundances for tHCT6 which is consistent with the experimentally measured abundances (Fig 6G). For pHCT6 an increase up to ~280% of wildtype levels was experimentally measured (Fig 6H). Our model was consistent with these results, estimating an increase up to ~265% of wildtype levels. Scenario 2 of the old model did not capture this increase in pHCT6, estimating wildtype levels for all three lines.

Overall, our model captured the increase from wildtype in all three of the proteins, while neither scenario of the old model captured the increase in pC3H3 and pHCT6. Scenario 2 of the old model estimated the increase in pPAL2 similar to the estimates from our model. The estimates from our model were consistent with the experimental tC3H3 and tHCT6 which were measured to remain around wildtype levels, and the measured increase in tPAL2.

### *PtrCAD1* and *PtrCAD2* knockdowns

Three experimental lines were analyzed where *PtrCAD1* and *PtrCAD2* were knocked down (Fig 7). From the differential abundance analysis, there were 18 transcripts and 12 proteins of untargeted genes that showed significant changes in abundance in at least one of the experimental lines (Figs 1C, 7A). We selected the *Ptr4CL3*, *PtrC4H1*, and *PtrCAld5H1* transcripts and proteins to compare the simulated results from our model with scenarios 1 and 2 of the old model. Fig 7B shows the amount that tCAD1 and tCAD2 were knocked down in the three experimental lines. For all three of these lines, tCAD1 was knocked down to ~5% of wildtype levels while tCAD2 ranged from no change from wildtype to ~25% of wildtype. These tCAD1 and tCAD2 abundances were used to emulate these knockdown experiments in our model and scenario 1 of the old model. For scenario 2 of the old model, measurements from all of the monolignol transcripts were used.

**Fig 7.**
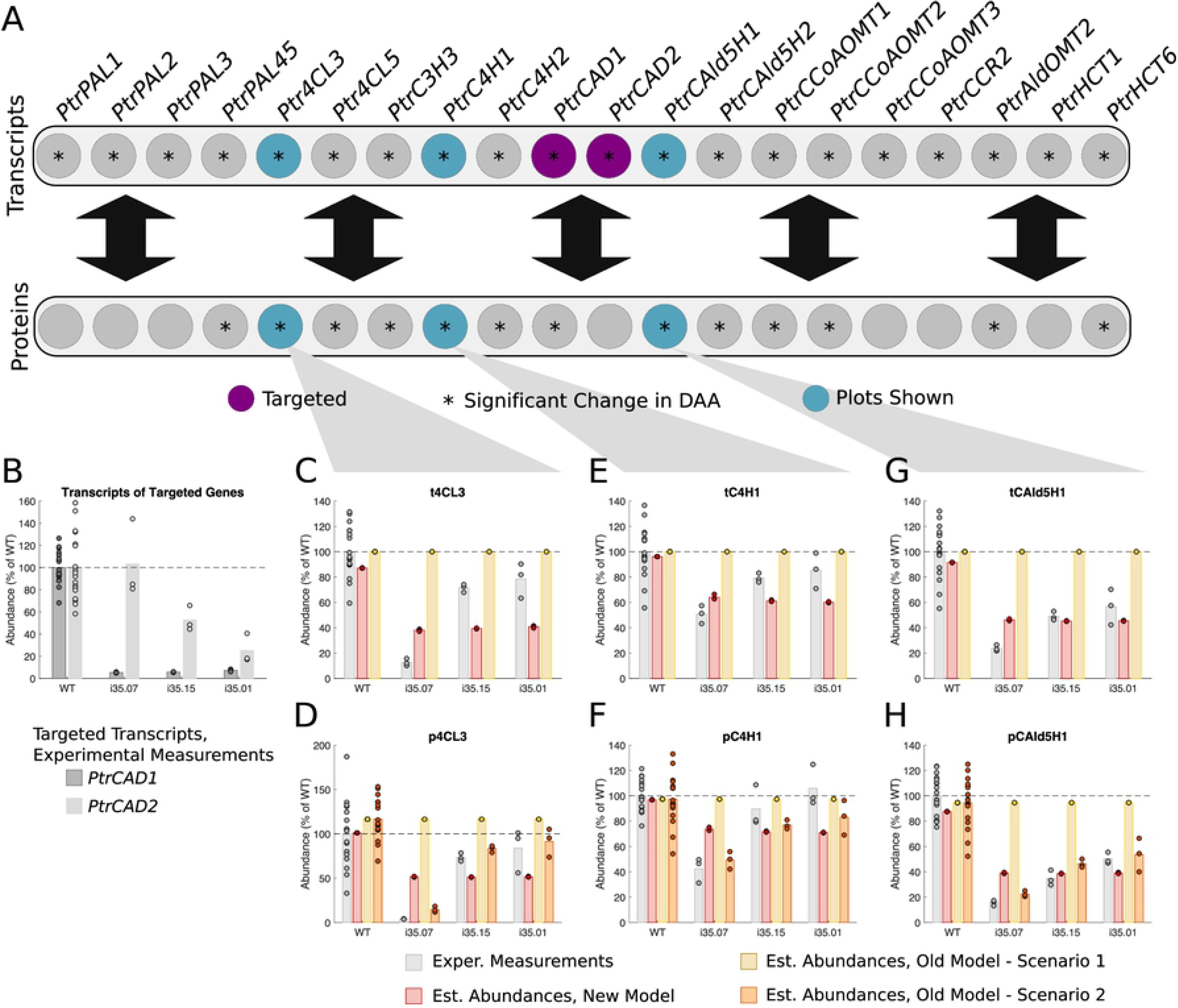
Experimental and estimated abundances of untargeted monolignol gene transcripts and proteins under *PtrCAD1* and *PtrCAD2* knockdowns. (A) Diagram showing targeted monolignol gene transcripts (purple), the transcripts and proteins that were found to have a significant change in abundance in at least one of the experimental lines (∗). (B) Level of knockdown of the targeted gene transcripts across the experimental lines. Experimental and estimated untargeted monolignol gene transcript and protein abundances for (C) t4CL3, (D) p4CL3, (E) tC4H1, (F) pC4H1, (G) tCAld5H1, and (H) pCAld5H1.

t4CL3 was experimentally measured in the range of ~80% to ~15% of wildtype levels (Fig 7C) and p4CL3 was experimentally measured in the range of ~85% to ~5% of wildtype levels (Fig 7D). Our model estimated a decrease to ~40% of wildtype levels for t4CL3 and a decrease to ~50% of wildtype levels for p4CL3 for all three lines, roughly consistent with the experimental measurements, though they do not capture the variation across the three lines. Scenario 2 of the old model estimated abundances of p4CL3 ranging from ~90% to ~15% of wildtype which is also consistent with the experimentally measured abundances.

For tC4H1 a decrease in abundance was experimentally measured ranging from ~85% to ~50% of wildtype levels (Fig 7E). A decrease in pC4H1 was experimentally measured ranging from ~100% to ~40% of wildtype levels (Fig 7F). Our model estimated a decrease to ~60% of wildtype levels for tC4H1 and a decrease to ~70% of wildtype levels for pC4H1 for all three lines, roughly consistent with the experimental measurements, though again, they do not capture the variation across the three lines. Scenario 2 of the old model estimated a decrease in the abundances of pC4H1 ranging from ~80% to ~50% of wildtype levels which are also consistent with the experimentally measured abundances.

A decrease in the abundance tCAld5H1 was experimentally measured ranging from ~60% to ~20% of wildtype levels (Fig 7G), and a decrease in pCAld5H1 was experimentally measured ranging from ~50% to ~15% of wildtype levels (Fig 7H). Our model estimated a decrease to ~45% of wildtype levels for tCAld5H1 and a decrease to ~40% of wildtype levels for pCAld5H1 for all three lines, consistent with the experimental measurements. Scenario 2 of the old model estimated decreases in pCAld5H1 ranging from ~55% to ~20% of wildtype levels, also consistent with the experimentally measured abundances.

Overall, scenario 2 of the old model did the best at estimating all three of the proteins because the decrease was captured in the transcript abundances. However, our model still captured the decrease from wildtype in both the transcripts and proteins despite only using the *PtrCAD1* and *PtrCAD2* transcript abundances as inputs to the model. The estimates from our model for the transcripts and proteins are very similar across the three experimental lines. This is due to the sparse maximum likelihood algorithm identifying *PtrCAD1*, which was knocked down similarly for all three lines, as a stronger influence on the other transcripts and proteins than *PtrCAD2*.

### *Ptr4CL3* and *Ptr4CL5* knockdowns

Three experimental lines were analyzed where *Ptr4CL3* and *Ptr4CL5* were knocked down (Fig 8). The differential abundance analysis identified 18 transcripts and 18 proteins of untargeted monolignol genes that showed significant changes in abundance in at least one of the experimental lines (Figs 1D, 8A). We selected the *PtrCAld5H2*, *PtrCCoAOMT3*, and *PtrHCT1* transcripts and proteins to compare the simulated results from our model with scenarios 1 and 2 of the old model. Fig 8B shows the different levels that t4CL3 and t4CL5 were knocked down for the three experimental lines. For all three of the lines, the transcripts were knocked down to around the same levels, ~5%-10% of wildtype levels. These t4CL3 and t4CL5 abundances were used to emulate these knockdown experiments in our model and scenario 1 of the old model. For scenario 2 of the old model, measurements from all of the monolignol transcripts were used.

**Fig 8.**
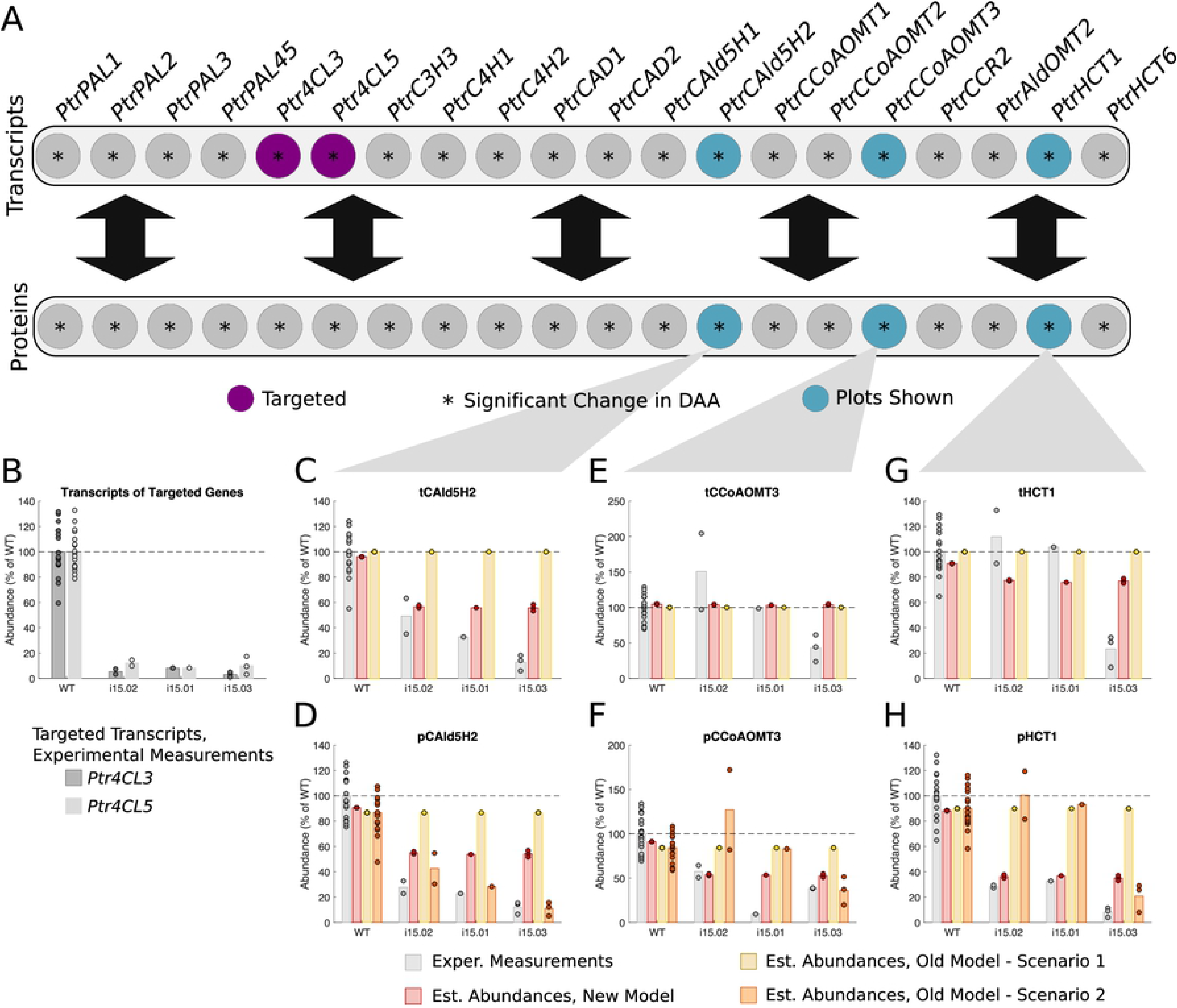
Experimental and estimated abundances of untargeted monolignol gene transcripts and proteins under *Ptr4CL3* and *Ptr4CL5* knockdowns. (A) Diagram showing targeted monolignol gene transcripts (purple), the transcripts and proteins that were found to have a significant change in abundance in at least one of the experimental lines (∗). (B) Level of knockdown of the targeted gene transcripts across the experimental lines. Experimental and estimated untargeted monolignol gene transcript and protein abundances for (C) tCAld5H2, (D) pCAld5H2, (E) tCCoAOMT3, (F) pCCoAOMT3, (G) tHCT1, and (H) pHCT1.

A decrease in abundance was experimentally measured in all three lines of tCAld5H2 ranging from ~50% to ~10% of wildtype levels (Fig 8C), and in pCAld5H2 ranging from ~30% to ~10% of wildtype levels (Fig 8D). Our model estimated a decrease to ~55% of wildtype levels for both tCAld5H2 and pCAld5H2 for all three lines. The decrease from wildtype in the estimated abundances is consistent with the experimental measurements, though the estimates from our model are not as low as the experimental values. Scenario 2 of the old model estimated a decrease in pCAld5H2 ranging from ~40% to ~10% of wildtype levels which is consistent with the experimentally measured abundances.

There was a large amount of variation in the experimentally measured tCCoAOMT3 abundances across the three lines, ranging from an average of ~150% of wildtype to ~45% of wildtype levels (Fig 8E). For pCCoAOMT3 a decrease in abundance was measured ranging from ~55% to ~10% of wildtype levels (Fig 8F). Our model estimated wildtype levels for tCCoAOMT3 and a decrease to ~55% of wildtype levels for pCCoAOMT3, consistent with the experimentally measured pCCoAOMT3. Due to the wide range in the measured transcript abundances, scenario 2 of the old model estimates protein abundances ranging from ~130% of wildtype to ~40% of wildtype levels. The estimates from scenario 2 of the old model for line i15-03 are consistent with the experimental measurements from that line, but its estimates from the other two lines, i15-02 and i15-01, are not consistent with the experimental measurements.

For two of the experimental lines, i15-02 and i15-01, tHCT1 was experimentally measured to be around wildtype levels. For the third line, i15-03, a decrease to ~25% of wildtype levels was measured (Fig 8G). However, a decrease in abundance was experimentally measured for pHCT1 in all three lines to ~30% of wildtype levels for lines i15-02 and i15-01 and to ~10% of wildtype levels for line i15-03 (Fig 8H). Our model estimated a slight decrease in tHCT1 to ~80% of wildtype levels for all three lines, and a decrease in pHCT1 to ~40% of wildtype levels in all three lines. The estimates for tHCT1 are roughly consistent with the experimental measurements for lines i15-02 and i15-01, but not for i15-03 which was much lower. The estimates for pHCT1 are consistent with the experimentally measured abundances for pHCT1 for all three lines. Because a decrease in tHCT1 abundance was only measured in line i15-03, scenario 2 of the old model estimated a decrease in pHCT1 only for that line, to ~20% of wildtype levels, consistent with the experimental measurements for that line. However, for the other two lines i15-02 and i15-01, scenario 2 of the old model estimated wildtype levels which are not consistent with the experimentally measured abundances from those two lines.

Scenario 2 of the old model did the best at estimating the *PtrCAld5H2* protein, but only estimated a decrease in the *PtrCCoAOMT3* and *PtrHCT1* proteins for the third line, i15-03. Our model, however, estimated a decrease in the abundances of all three proteins for all three of the lines. Our model also captured the decrease in tCAld5H2, and its estimates for tCCoAOMT3 and tHCT1 are reasonable considering the range of the measured abundances across the three lines.

### *PtrCCoAOMT3* knockdowns

Three experimental lines were analyzed where *PtrCCoAOMT3* was knocked down (Fig 9). The differential abundance analysis identified 3 transcripts and 16 proteins of untargeted monolignol genes that had significant changes in abundance in at least one of the experimental lines (Figs 1E, 9A). We selected the *Ptr4CL3*, *PtrCAD1*, and *PtrHCT1* transcripts and proteins to compare the simulated results from our model with scenarios 1 and 2 of the old model. Fig 9B shows the range that tCCoAOMT3 was knocked down over the 3 experimental lines. In the first line, i21-03, tCCoAOMT3 was not knocked down from wildtype. In the other two lines it was knocked down to ~20% of wildtype levels. These tCCoAOMT3 abundances were used to emulate these knockdown experiments in our model and scenario 1 of the old model. For scenario 2 of the old model, measurements from all of the monolignol transcripts were used.

**Fig 9.**
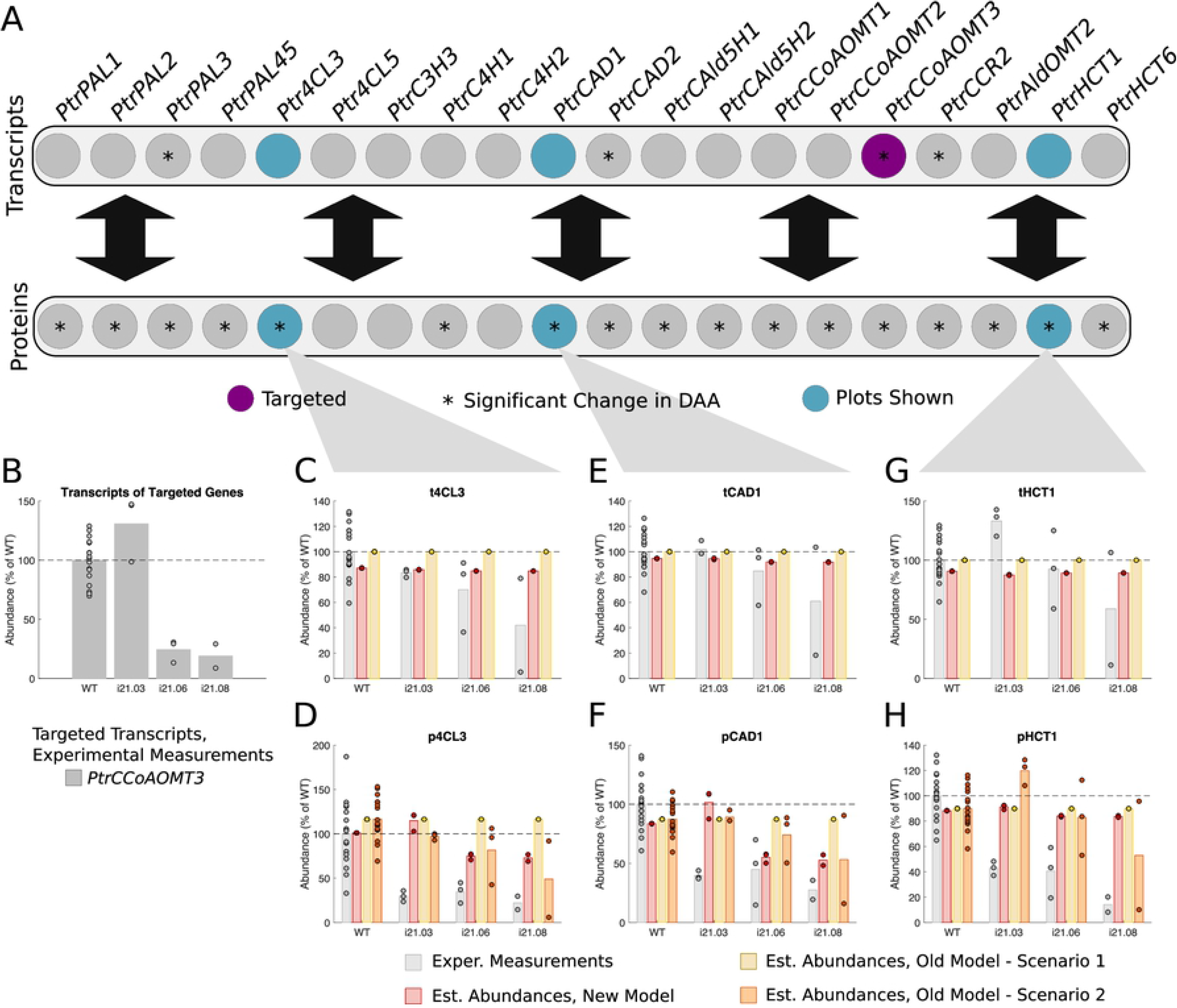
Experimental and estimated abundances of untargeted monolignol gene transcripts and proteins under *PtrCCoAOMT3* knockdowns. (A) Diagram showing targeted monolignol gene transcripts (purple), the transcripts and proteins that were found to have a significant change in abundance in at least one of the experimental lines (∗). (B) Level of knockdown of the targeted gene transcripts across the experimental lines. Experimental and estimated untargeted monolignol gene transcript and protein abundances for (C) t4CL3, (D) p4CL3, (E) tCAD1, (F) pCAD1, (G) tHCT1, and (H) pHCT1.

A decrease in abundance was experimentally measured for t4CL3 ranging from wildtype levels to ~40% of wildtype levels (Fig 9C). However there is a large amount of variation between replicates, especially for lines i21-06 and i21-08. Our model did not estimate any change from wildtype levels for all three lines. A decrease in p4CL3 was experimentally measured ranging from ~35% to ~20% of wildtype levels (Fig 9D). Our model only estimated a decrease to ~75% of wildtype levels and scenario 2 of the old model estimated a decrease ranging from no change from wildtype to ~50% of wildtype, neither of which are very consistent with the decrease that was experimentally measured.

A decrease in abundance was experimentally measured for tCAD1 ranging from wildtype levels to ~60% of wildtype levels (Fig 9E). However there is, again, a large amount of variation between replicates, especially for lines i21-06 and i21-08. Our model did not estimate any change from wildtype levels for all three lines. A decrease in pCAD1 was experimentally measured ranging from ~45% to ~25% of wildtype levels (Fig 9F). For the two lines where tCCoAOMT3 was decreased, i21-06 and i21-08, our model estimated a decrease in pCAD1 to ~55% of wildtype levels, consistent with the expermental values. For the same lines, scenario 2 of the old model estimated a decrease in pCAD1 ranging from ~75% to ~55% of wildtype levels.

A decrease in abundance was experimentally measured for tHCT1 ranging from ~120% to ~50% of wildtype levels (Fig 9G). Again, there is a large amount of variation between replicates, especially for lines i21-06 and i21-08. Our model did not estimate any change from wildtype levels for tHCT1 for all three lines. A decrease in pHCT1 was experimentally measured ranging from ~45% to ~15% of wildtype levels (Fig 9H). Our model did not estimate any change from wildtype levels for all three lines, while estimates from scenario 2 of the old model ranged from ~120% to ~50% of wildtype levels, neither of which are very consistent with the decrease that was experimentally measured.

Overall, neither our model, nor the old model, did a good job at estimating the experimentally observed changes for the *Ptr4CL3* and *PtrHCT1* transcripts and proteins. However, our model was able to better capture the decrease in pCAD1 than scenario 2 of the old model.

## Discussion

Significant work has been done in recent years to understand the transcriptional regulation of monolignol biosynthesis and wood formation [12, 41, 42]. Chen et al., [12] recently constructed a heirarchical transcriptional regulatory network for wood formation in *P. trichocarpa*. They identified 7 transcription factors (TFs) that regulated 10 of the monolignol specific genes: *PtrPAL2*, *PtrCCoAOMT1*, *PtrCCoAOMT2*, *PtrCAld5H1*, *PtrCAld5H2*, *PtrAldOMT2*, *PtrCAD1*, *PtrHCT1*, *PtrHCT6*, and *PtrC4H1* [12]. In the *PtrCAD1* and *PtrCAD2* transgenics (Fig 1C) we found these 10 transcripts, among others, to be differentially expressed. Many of the TFs that Chen et al., identified as regulators of these genes were also found to be differentially expressed in these transgenics (Fig S5), further supporting that the cross-influences impacting the abundances of these transcripts are occuring through TF regulation.

In addition to changes in transcript abundance, we also observed several cases where monolignol protein abundances were significantly altered when their transcripts were not. This behavior has previously been observed in secondary cell wall proteins of *Arabidopsis* [37, 38] and in tobacco during cell differentiation [39]. Compared to transcriptional regulation, less is known about the role of post-transcriptional and post-translational regulatory mechanisms on monolignol biosynthesis. Phosphorylation of the *PtrPAL* protein was proposed for monolignol biosynthesis over two decades ago, though the role of this phosphorylation is unknown [43, 44]. Wang et al., [15] characterized the phosphorylation of the *PtrAldOMT2* protein in *P. trichocarpa*. This post-translational modification was found to impact the activity of the *PtrAldOMT2* protein but not its abundance. Loziuk et al., identified 12 monolignol proteins that contain motifs for potential glycosylation in *P. trichocarpa* [16]. The proteins they identified include pPAL1, pPAL3, pPAL4, pPAL5, pC3H3, p4CL3, pCAD2, pCAld5H2, pCCoAOMT1, pCCoAOMT2, pCCR, and pHCT1. Like phosphorylation, glycosylation can regulate protein localization, functional activity, ability to form multienzyme complexes, and stability [16].

Glycosylation could explain some of the behavior we observed in the protein abundance data. In the *PtrC3H3*, *PtrC4H1*, and *PtrC4H2* knockdowns (Fig 1A) and the *PtrCAld5H1* and *PtrCAld5H2* knockdowns (Fig 1B) we observed significant changes in the *PtrPAL*, *PtrHCT*, *PtrCCoAOMT2*, *PtrCAld5H*, *PtrC3H3*, and *Ptr4CL* proteins. At least one protein in each of those families was found to have glycosylation motifs [16]. The *PtrHCT* proteins, particularly, had significant changes in their protein abundances, which were not observed in their transcripts across multiple transgenic knockdowns (Fig 1A,B,D,E, Fig S4A,B), or where their transcripts were differentially expressed but the proteins were not significantly different from wildtype (Fig 1C, Fig S2B). Further, there appear to be relationships among the *PtrHCT*, *Ptr4CL*, *PtrCCoAOMT3*, and the *PtrCAD1* proteins (Fig 1D,E, Fig S4A,B), with reciprocal indirect influences between the *Ptr4CL* and *PtrCCoAOMT3* proteins, suggesting a potential feedback mechanism. The sparse maximum likelihood estimator detected several connections among these proteins, including positive influences from p4CL3, p4CL5, pCAD1 and pHCT1 on pCCoAOMT3, from p4CL5, pCCoAOMT3, and pHCT1 on pCAD1, from the p4CLs and pCCoAOMT3 on the pHCTs, and from pCCoAOMT3 and the pHCTs on the p4CLs (Fig 4A). Further experiments are needed to identify the specific regulatory mechanisms that are responsible for these cross-influences.

We used the connections identified by the sparse maximum likelihood estimator to define our new transcript-protein model for monolignol biosynthesis. Using this model, we emulated the 225 wildtype and transgenic knockdown experiments using only the measured transcript abundances from the targeted monolignol genes as an input and estimating the abundances of the other, untargeted, transcripts and proteins. We compared these estimates to those found using the old model [4], which assumes the protein abundances are linearly proportional to the transcript abundance of the same monolignol gene. We performed a 10×10-fold cross-validation and compared the resulting RMSE distributions from the old model and our new model. The mean RMSEs for 14 of the 20 transcripts and 11 of the 20 proteins were found to be statistically lower in our new model than the old model. We then simulated the transgenic experiments from our differential abundance analysis using our model and scenarios 1 and 2 of the old model, and compared the estimated transcript and protein abundances of selected untargeted genes of interest. As expected, scenario 2 of the old model, which uses the full transcript abundance profiles, did the best at estimating the proteins whose abundance levels tracked the abundance levels of its transcripts, such as *Ptr4CL3*, *PtrC4H1*, and *PtrCAld5H1* in the *PtrCAD1* and *PtrCAD2* knockdown experiments (Fig 7C-H), and *PtrCAld5H2* in the *Ptr4CL3* and *Ptr4CL5* knockdowns (Fig 8C-D). However, using only the targeted *PtrCAD1* and *PtrCAD2* or *Ptr4CL3* and *Ptr4CL5* transcripts respectively, our model was still able to estimate the decreases in both the transcripts and proteins for all four of these genes. Additionally, our model was able to capture several changes in protein abundances that the old model was not, including *Ptr4CL5*, *PtrCAld5H2*, and *PtrHCT1* in the *PtrC3H3*, *PtrC4H1*, and *PtrC4H2* knockdowns; *PtrC3H3* and *PtrHCT6* in the *PtrCAld5H1* and *PtrCAld5H2* knockdowns; and *PtrCCoAOMT3* and *PtrHCT1* in the *Ptr4CL3* and *Ptr4CL5* knockdowns.

Neither model was able to estimate the changes in abundance of the *Ptr4CL3* and *PtrHCT1* proteins in the *PtrCCoAOMT3* transgenics. Our model includes relationships from pCCoAOMT3 to p4CL3 and pHCT1. Despite this, our model does not capture the size of the decrease in the abundances of these proteins. One explanation for why the extent of these regulatory influences are not captured in our simulations could be due to constraining the regulatory influences to additive linear relationships. Some of the shortcomings of an additive linear model include not allowing for nonlinear relationships and not being able to capture synergistic influence behaviors (i.e., when multiple components are needed to see an effect).

The monolignol proteins are the driving forces in the biosynthesis pathway, so being able to accurately understand and estimate how they change under different combinations and degrees of targeted genetic modifications is important for the accuracy of predictive models. Regulatory influences that occur after transcription appear in the monolignol data of stem differentiated xylem tissue in *P. trichocarpa*, and we have developed a computational model that incorporates influences on both the monolignol transcripts and proteins. We have demonstrated specific examples where our model produces better estimates of experimental monolignol gene proteins than the old model when both models use only the targeted monolignol transcript abundances as input. In several cases our model, using only the targeted transcript abundances, produced better estimates than scenario 2 of the old model where all of the experimental transcript abundances were used. By incorporating these indirect regulatory influences, we believe our model has improved ability to explore the cascaded impact of genetic modifications on resulting lignin and wood characteristics. Future work will evaluate how our model performs on independent data, incorporate the model into the multi-scale model in [4], and use the multi-scale model to explore the possible changes in lignin and wood characteristics under combinations of lignin gene modifications.

## Methods

### Monolignol transcript-protein model

The multi-scale lignin biosynthesis model presented in [4] spans multiple biological layers from the genome to observed lignin and wood physical and chemical traits. However, that model [4] makes the simplifying assumption that each monolignol gene’s protein abundance is dependent only on its transcript abundance. This does not reflect any changes that are observed in the abundance of the non-targeted genes. Here, we present a new model that incorporates the observed influences that estimate the production of untargeted monoligninol transcripts and proteins. The code associated with this model can be found at https://github.ncsu.edu/mlmatth2/Monolignol-Cross-Regulation-Model.

Because we are interested in identifying regulatory influences at not only the transcriptional level, but also the translational level, we combined the two datasets, such that we are now looking at each of the 20 transcripts and 20 proteins as 40 total variables in our model.

### Model development

The goal of the model development is to find the underlying influences on each monolignol gene product (its transcripts and proteins) when the expression of other monoligninol genes are modified. We describe each transcript and protein as a linear combination of the other transcripts and proteins as shown in Eq (1).

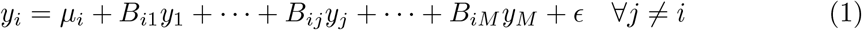

Where *y*_*i*_ is the abundance of the *i*^*th*^ gene product, and we have *M* total gene products (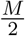 transcripts and 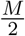 proteins). *B*_*ij*_ is a constant term that reflects the influence of gene product *j* on gene product *i*, *μ*_*i*_ is a constant that represents the portion of *y*_*i*_ that is not described by the other lignin gene products, and *ϵ* is the error. The influences described by *B*_*ij*_ should be consistent across multiple experiments, so we can describe Eq (1) over a collection of experiments as shown in Eq (2).

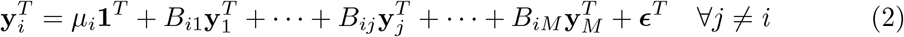

Where 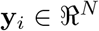 is the abundances of *i*^*th*^ gene product over *N* experiments. We can combine this into one model for all the transcripts and proteins as shown in Eq (3).

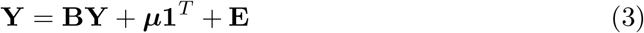

Where 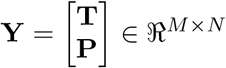 is a matrix composed of the abundances for the 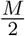 transcripts (**T**) and associated 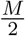 proteins (**P**) for each of the *N* experiments. 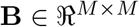 is the collection of influence terms *B*_*ij*_. Because each *y*_*i*_ is a function of the other gene products *y*_*j*_ ∀*j* ≠ *i*, the diagonal elements of **B**, *B*_*ii*_ = 0 ∀*i*. Additionally, we also enforce a constraint that a transcript cannot be influenced by its associated protein 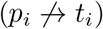. 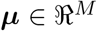 is a vector containing a constant term for each gene product, and 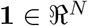 is a vector of all ones. **E** = [***ϵ***_1_ ***ϵ***_2_ · · · ***ϵ***_*N*_ l represents the error where 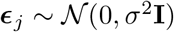 and is considered independent and identically distributed.

We used a sparse maximum likelihood (SML) estimator [29] adjusted for our model and data structure (S1 Text) to solve for **B** and ***μ***. SML adds an *l*_1_-norm regularization term to the maximum likelihood, encouraging elements of **B** to be zero if they are not sufficiently useful to describing **Y**. A coordinate-ascent algorithm is used, allowing us to solve for the influences defined in **B** and ***μ*** on a row-by-row basis as described in Eq (2). This allows us to control which experiments are used to solve for the *i*^th^ row of **B** and ***μ***, 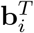 and *μ*_*i*_ respectively. This is important because we do not want to include the experiments where component *i* was targeted. In those experiments, an outside influence that is not included in the model is impacting its abundance. Only transcripts were considered to be targets at this stage, as those are what is directly modified in the knockdown experiments. See S1 Text for more details on the model development and SML approach.

### Estimating monolignol transcripts and proteins

We can use the influences **B** and ***μ*** solved for in the model development stage and Eq (4) to estimate how knocking down a single or combination of monolignol genes alters the abundances of the untargeted monolignol transcripts and proteins.

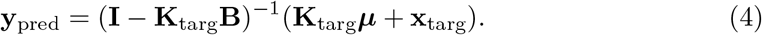

We set the abundance of our targeted components to the desired knocked down amount using the vector 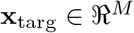, and remove the model influences that would alter these set abundances using 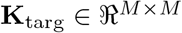. Where **X**_targ_ = Σ_*i*∊targ_*x*_*i*_**e**_*i*_ and 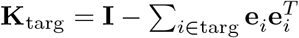. **e**_*i*_ is the *i*^th^ unit vector. This configuration allows us to set the targeted monolignol gene components to a desired value while keeping the relationships that influence the untargeted monolignol transcripts and proteins.

A drawback of using the additive linear model to describe both the monolignol transcripts and proteins, is that a complete knockout of a targeted transcript may not result in our model estimating its protein to be completely knocked out as well. This presents an issue if the goal is to examine the impact of complete knockouts of targeted monolignol genes. To get around this issue, we assume that the targeted change in a transcript results in a proportional change to its protein abundance. For example, if we want to see what happens when we knock transcript 1, *t*_1_ down to 10% of its wildtype abundance, then 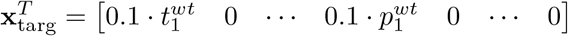 and 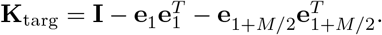

### Differential abundance analysis

We performed the differential abundance analysis for the monolignol gene transcripts [4] using the R package DESeq2 [45] for each batch individually using the RNA-seq libraries available under GEO accession number GSE78953. The proteomics data [4] was log2 transformed and the limma package [46, 47] was used for each batch to identify significant differential abundance [48]. The proteomics data set is available on CyVerse (http://mirrors.iplantcollaborative.org/browse/iplant/home/shared/LigninSystesmDB).

### Missing data imputation

In the proteomics data set, 83 out of the 4500 proteins measured (1.8%) could not be quantified. We employed a series of rules to estimate these missing values: 1) If the protein was successfully measured for at least one other replicate in the same line, then the missing value was replaced with the average abundance of the protein from the other replicates of that line. This accounted for 42 of the missing values. 2) If a protein was not quantified for all replicates of an experimental line, then 2a) if the missing value is for a protein associated with the monolignol gene targeted for knockdown, we replaced the missing value with the fraction of its average wildtype abundance that its associated transcript was knocked down. For example, if the associated transcript was knocked down to 10% of its average wildtype value, then the missing protein value was replaced with 10% of its average wildtype value. This accounted for 30 of the missing values. 2b) The remaining missing values were replaced with the average wildtype value of that protein. This accounted for 11 of the missing values.

## Supporting information

### S1 Text. Supporting information

**S1 Fig. Monolignol gene transcript and protein differential abundance (cont.).** (A) *PtrPAL1* knockdown experiments (Construct a1). (B) *PtrPAL2*, *PtrPAL4*, and *PtrPAL5* knockdown experiments (Construct i7). (C) *PtrPAL4* knockdown experiments (Construct a3). (D) *PtrPAL5* knockdown experiments (Construct a4). (E) *PtrPAL2* knockdown experiments (Construct a5). (F) *PtrPAL1* and *PtrPAL3* knockdown experiments (Construct i6). Gray boxes are due to missing data. Rows are the monolignol gene names, with the targeted genes for each experiment in purple. Columns are the experimental lines. ∗ indicates p_adj_<0.05.

**S2 Fig. Monolignol gene transcript and protein differential abundance (cont.).** (A) *PtrPAL1*-*PtrPAL5* knockdown experiments (Construct i8). (B) *PtrC3H3* knockdown experiments (Construct i20). (C) *PtrCAD1* knockdown experiments (Construct i33). (D) *PtrC4H2* knockdown experiments (Construct a9). (E) *PtrC4H1* knockdown experiments (Construct a10). (F) *PtrCCR2* knockdown experiments (Construct i26). Gray boxes are due to missing data. Rows are the monolignol gene names, with the targeted genes for each experiment in purple. Columns are the experimental lines. ∗ indicates p_adj_<0.05.

**S3 Fig. Monolignol gene transcript and protein differential abundance (cont.).** (A) *PtrHCT1* knockdown experiments (Construct a17). (B) *PtrHCT6* knockdown experiments (Construct a18). (C) *PtrCCoAOMT1* knockdown experiments (Construct a22). (D) *PtrCAld5H1* knockdown experiments (Construct a27). (E) *PtrCAld5H2* knockdown experiments (Construct a28). (F) *PtrHCT1* and *PtrHCT6* knockdown experiments (Construct i19). Gray boxes are due to missing data. Rows are the monolignol gene names, with the targeted genes for each experiment in purple. Columns are the experimental lines. ∗ indicates p_adj_<0.05.

**S4 Fig. Monolignol gene transcript and protein differential abundance (cont.).** *Ptr4CL3* knockdown experiments (Construct a12). (B) *Ptr4CL5* knockdown experiments (Construct a13). (C) *PtrCCoAOMT1* and *PtrCCoAOMT2* knockdown experiments (Construct i24). (D) *PtrAldOMT2* knockdown experiments (Construct i30). Gray boxes are due to missing data. Rows are the monolignol gene names, with the targeted genes for each experiment in purple. Columns are the experimental lines. ∗ indicates p_adj_<0.05.

**S5 Fig. Transcription factor expression in *PtrCAD1* and *PtrCAD2* knockdowns.** Rows are the TFs identified in [12] that regulate the monolignol genes. Columns are the experimental lines. ∗ indicates p_adj_<0.05.

**S1 Table. Table of relationships identified using SML approach.**

## Acknowledgments

This work was supported by NSF Grant DBI-0922391 and a National Physical Science Consortium (NPSC) Graduate Fellowship. We also thank David C. Muddimann for his work quantifying the proteomics used in this manuscript.

